# The scientific basis for currently proposed management options for Hector’s and Maui dolphins: a critique

**DOI:** 10.1101/2020.05.15.098889

**Authors:** Elisabeth Slooten, Stephen Michael Dawson

## Abstract

New Zealand’s Ministry for Primary Industries (MPI) developed a risk analysis for Hector’s and Maui dolphins, in order to inform protection options being considered by the Ministers of Fisheries and Conservation. Unfortunately, MPI’s risk analysis combines several estimates that are biased, and the biases consistently act together to underestimate the level of bycatch and overestimate the species’ ability to sustain impacts. In essence, the approach uses abundance estimates that are likely biased high, multiplies them by a reproductive rate that has been arbitrarily raised, multiplied by an assumed figure for calf survival, to reach a number of dolphins that would be added each year if the population were to remain stable. From this number, they subtract their estimates of bycatch, which are almost certainly biased low. The remaining number of dolphins is apportioned a cause of death according to autopsy data from 55 Hector’s and Maui dolphins found dead on beaches. This is then compared to estimates of what level of takes would be sustainable, calculated using a formula that is not well understood and less conservative than the international standard (PBR, developed by the US National Marine Fisheries Service). This is a poor basis for rational management of an endangered, endemic marine mammal. A key problem with the MPI approach is that its definition of “risk” does not relate to the risk of population decline or extinction, and is inconsistent with the modern understanding of the behaviour of meta-populations. The approach defines risk as the likelihood of capture, which is apportioned to different areas according to fishing effort and the habitat model’s outputs for dolphin distribution. The protection options are therefore targeted where high densities of dolphins and high fishing effort coincide. Large populations are allocated the highest level of protection, while small populations remain poorly protected. This approach is likely to increase the risk of local extinctions, contractions of dolphin distribution, population fragmentation, loss of genetic variability and result in increased risk to the species as a whole. Many of these points were made by an International Expert Panel in 2018. MPI have failed to provide a list of recommendations from the Expert Panel Report with their responses (if any) to those recommendations. This standard step in scientific practice, following peer review (e.g. response to reviewers’ comments) has not been followed.

## INTRODUCTION

The management options presented in the Hector’s and Maui Dolphin Threat Management Plan (TMP 2019) are based on the spatial risk assessment discussed at the 2019 IWC meeting (Roberts et al. 2019). In developing the TMP, the Ministry for Primary Industries (MPI) approach has been to fill in gaps in sightings data and fisheries data with a complex spatial risk assessment. For reasons described below, this has failed to produce a scientifically robust basis for management decisions. An additional problem is that the management goals in the TMP are neither specific nor measurable.

In this contribution, Section A evaluates options presented in the Threat Management Plan (TMP 2019) for reducing bycatch of Hector’s and Maui dolphins. Section B presents an overview of the problems with the science underpinning these management options (Roberts et al. 2019). We find that flaws in the MPI assessment combine to create a biased assessment of risks. Hence the options in the TMP (2019) are not appropriately informed by robust science.

### A. OPTIONS FOR REDUCING BYCATCH PROPOSED IN THE THREAT MANAGEMENT PLAN

#### Maui dolphin

Following the extinction of baiji (Turvey et al. 2007) and the apparently imminent extinction of vaquita (Jaramillo-Legorreta et al. 2019), marine mammalogists have learned that Critically Endangered subspecies such as Maui dolphin need highly effective protection in order to recover to non-threatened status. For Maui dolphin, Such protection would be achieved by eliminating gillnetting and trawling, throughout their habitat, out to the 100m depth contour or 20 nautical miles offshore. This includes banning gillnetting within the harbours, which are part of Maui dolphin habitat (Rayment et al. 2011).

In contrast, the TMP’s most far-reaching option (Option 4, Figure 1) offers negligible extra protection south of New Plymout, a zone in which Maui dolphins were routinely present a few decades ago (Dawson et al 2001, McGrath 2020). It is not clear why less protection is proposed in areas with small populations of Maui (and Hector’s) dolphin. These are the populations at greatest risk, yet would remain poorly protected under Option 4.

**Figure 1.**
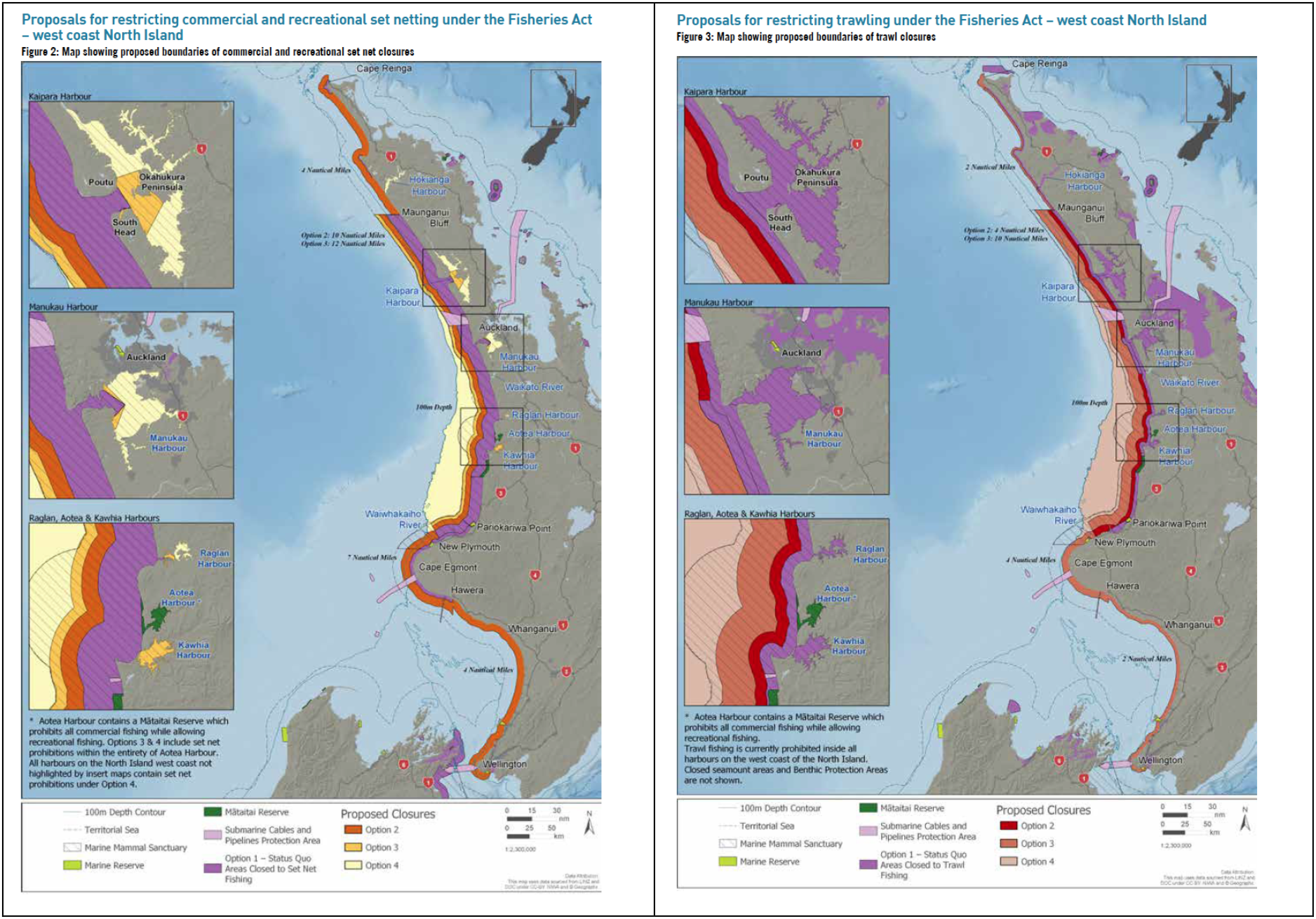
Protection options for Māui dolphins from gillnet (left) and trawl fisheries (right) in the Threat Management Plan.

The TMP outlines a Toxoplasmosis Action Plan as part of the Department of Conservation’s (DOC) response. While several Maui dolphins have died of toxoplasmosis (Roe et al. 2013), it is clear that the impact of this disease has been overstated in the TMP (see Section B below). Eliminating cats in the greater Auckland area would bring many general conservation benefits (e.g. for urban bird populations). However, public opposition is likely to render this unachieveable and it is not clear whether eliminating cats would materially benefit Maui dolphin. Focussing on wild cats, as has been suggested by DOC in public meetings, ignores the fact that abandoned domestic cats become wild cats. While a domestic population remains, there will be a ready source of recruits to the wild population. For these reasons, we see the currrent toxoplasmosis action plan as having a low chance of success. Further, it is critical to ensure that efforts to study and manage toxoplasmosis do not distract attention and resources from addressing direct impacts that can be readily managed – in particular fisheries bycatch.

#### Hector’s dolphin

Fundamental problems with the science underpinning the TMP process have resulted in management options that are poorly focussed and highly likely to have negative consequences for several local and regional populations. Option 3, the most effective option for Hector’s dolphin (Figure 2) involves extending existing protection only in Te Waewae Bay, the greater Banks Peninusla area, off Kaikoura and in Tasman and Golden Bays. Important regional populations off southeastern and northeastern coasts of the South Island have been ignored. These populations have had recent gillnet mortalities, and their low abundance renders them especially vulnerable to decline. None of the proposed options improve protection off the South Island west coast. Due to extremely low levels of observer coverage, quantitative information on bycatch in this area is lacking.

**Figure 2.**
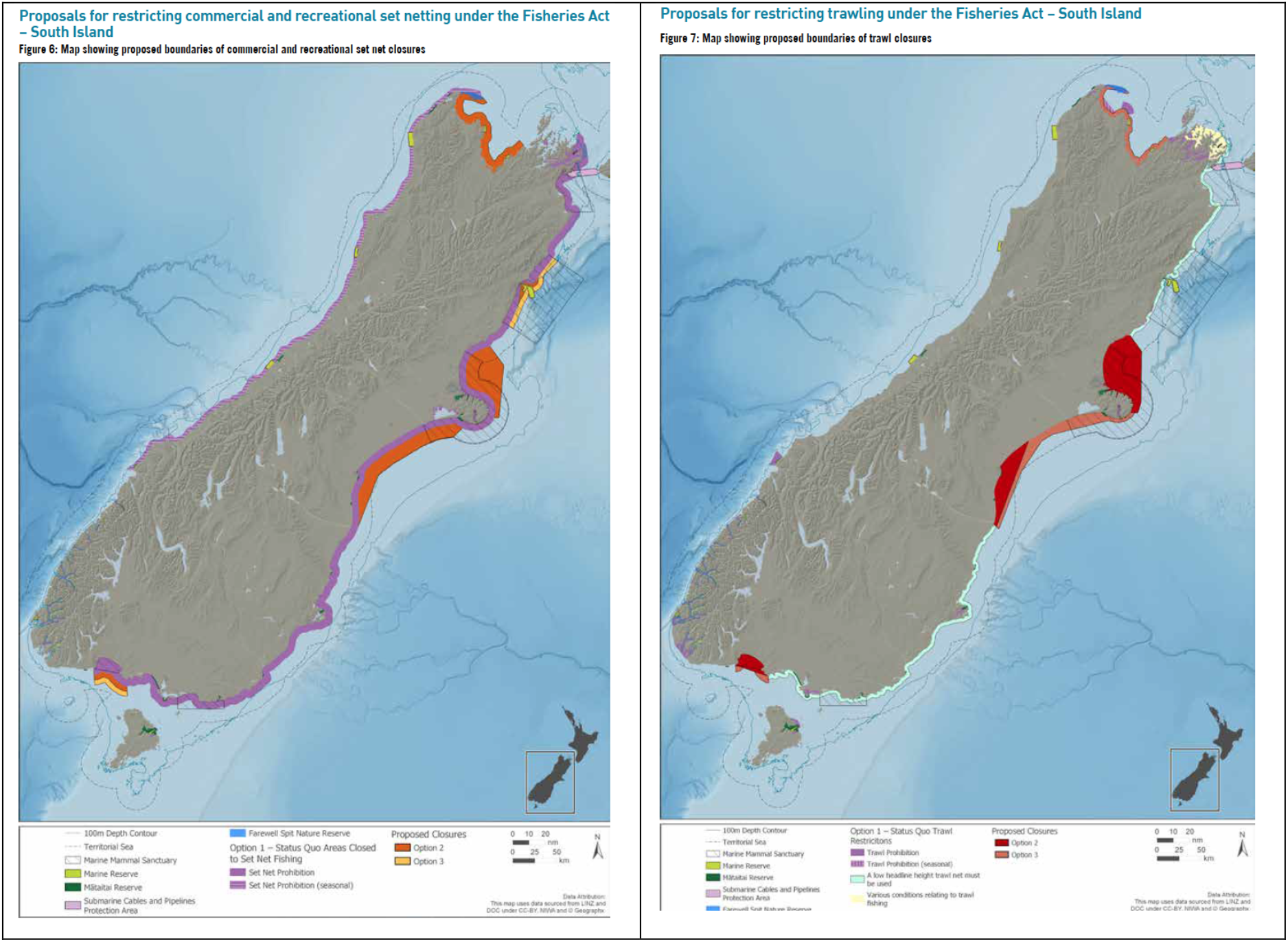
Protection options for Hector’s dolphins from gillnet (left) and trawl fisheries (right) in the Threat Management Plan.

The gap in protection off the south-east coastline of Banks Peninsula (Figures 3 and 4) is surprising. This is a well documented hotspot of Hector’s dolphin abundance (e.g. Rayment et al. 2010). This omission results directly from the habitat model that has been used to support the TMP (see Part B below) and the assumption that fishing effort will not be displaced in response to new protection measures. Past experience shows that fishing effort will be displaced to areas where fishing restrictions are less stringent; nearby at first, but ultimately venturing as far as fishers feel they need to. The proposed gap in protection is likely to become a focus of intense fishing activity. Because dolphin abundance there is high, this is likely to increase bycatch, and result in the area becoming a population sink. This would result in shifting, rather than solving the problem.

**Figure 3.**
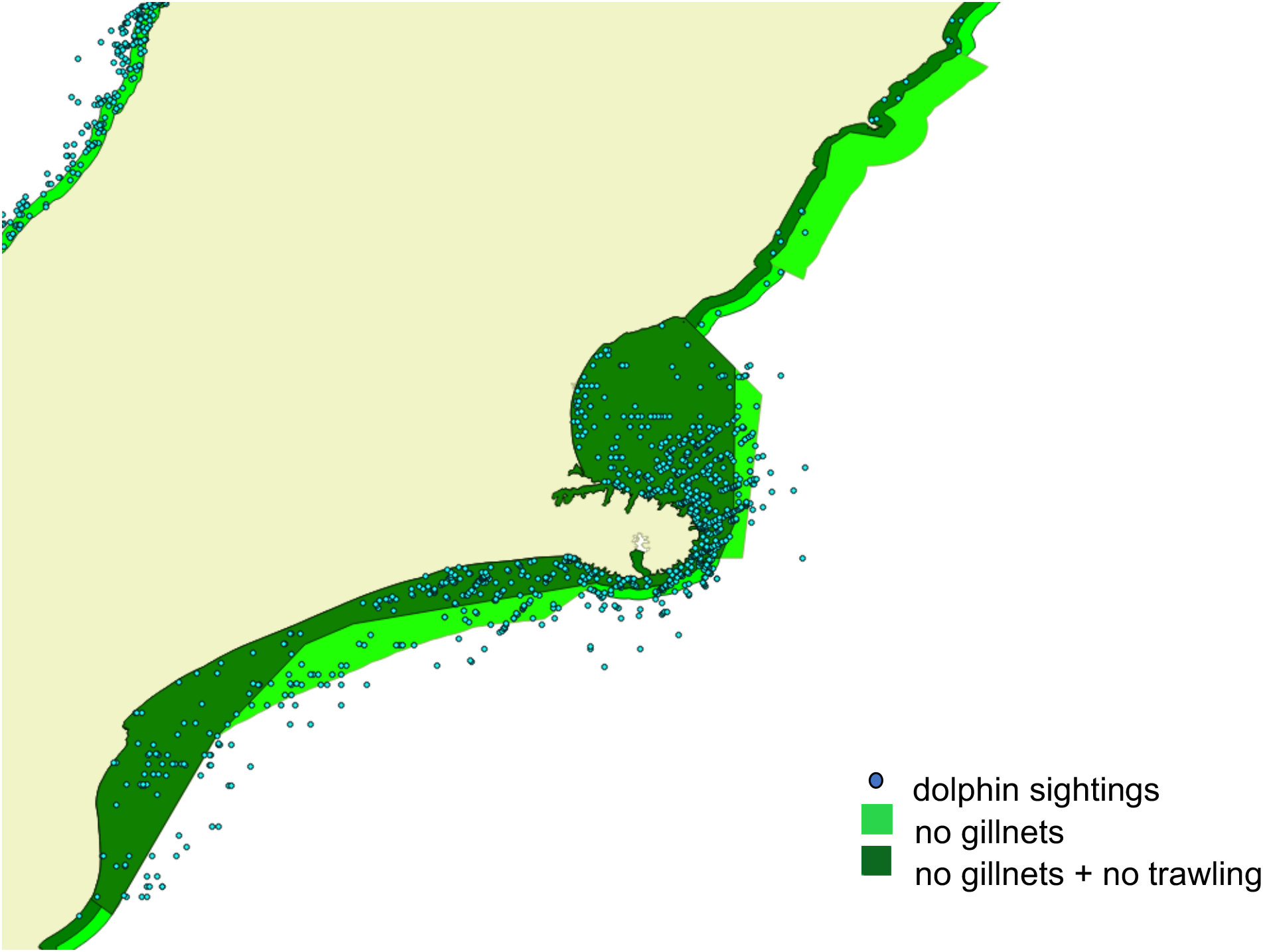
Poor fit between dolphin sightings and the proposed protection measures (Option 3), based on the MPI habitat model.

**Figure 4.**
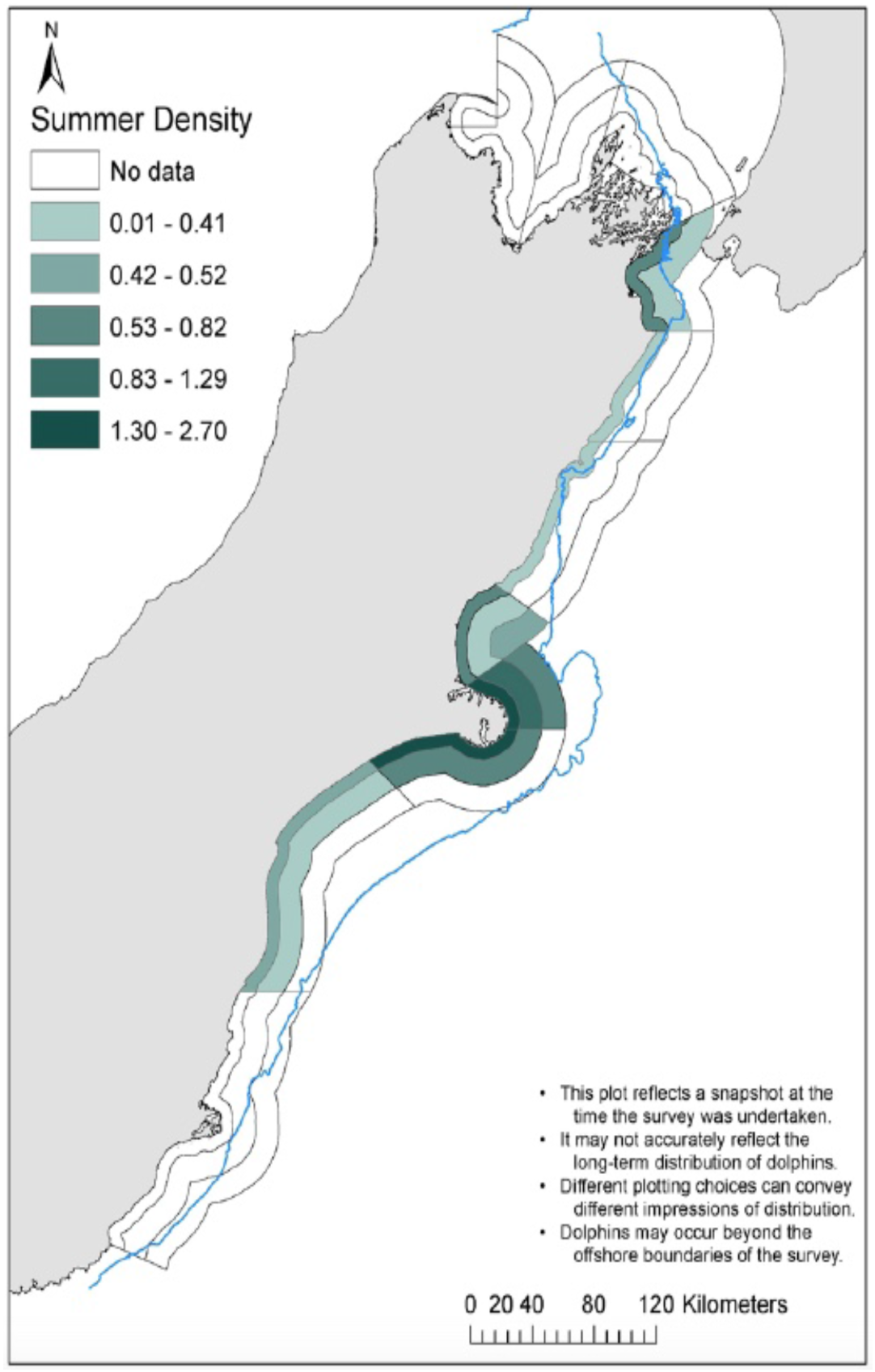
The latest survey results (MacKenzie and Clement 2014) indicate high dolphin densities in the area left out of the proposed protection measures on the southeastern side of Banks Peninsula (Figure 3).

Possibly the most serious oversight is the lack of attention given to small, local and regional populations (Taylor et al. 2018). Photo-ID and genetic studies have shown small populations, off the Catlins, Otago and in Cloudy Bay, to be resident, with reasonably precise mark-recapture estimates of abundance (Turek et al. 2013, Hamner et al. 2017, Harvey, unpub. data). These local populations connect the larger populations in Te WaeWae Bay and Canterbury, and provide a conduit for genetic and demographic connectivity. If they are permitted to decline to extinction, the current fragmentation of the species’ total population (Hamner et al. 2012) will increase, increasing risk to the species as a whole.

The distribution of fishing effort can be highly labile, particularly in the face of fisheries regulations. For this reason, efforts to solve bycatch problems based on the distribution of current or recent fishing effort are likely to compromise long-term effectiveness. When fishers shift, bycatch problems will likely to shift with them. This problem has been recognised for decades (e.g. Dawson 1991). The solution is to ensure there is effective protection where the dolphins are, rather than only where gillnet and trawl fisheries are currently operating. This strategy has no cost to fishers if they are not operating in a particular area currently, and protects dolphin populations from shifts in fishing effort, whether caused by regulation, stock depletion, or market pressure.

For the reasons outlined above, the protection measures proposed in the TMP are inadequate. Even the best of them (Option 3) is very unlikely to be effective in reducing bycatch. It is unfortunate that IWC and IUCN recommendations for protecting Maui and Hector’s dolphins were not included as options in the TMP when it went out for public consultation.

### B. PROBLEMS WITH THE SCIENCE SUPPORTING THE TMP

The Ministry for Primary Industries’ (MPI) threat assessment approach (Roberts et al. 2019) relies critically on information on dolphin distribution, population size, reproductive rate, fishing effort and the catch rate of dolphins per unit of fishing effort. This section briefly outlines the main problems with the approach taken, and the data inputs used in the risk assessment (Roberts et al. 2019) supporting the TMP.

#### 1. Population size

The population surveys carried out under contract for MPI have problems that have been pointed out in several peer reviews. The east coast South Island survey (MacKenzie and Clement 2014) in particular, is likely to be biased high, for reasons explained elsewhere (e.g. IWC 2016). It is treated at face value in the analyses supporting the TMP.

#### 2. Habitat model

The population surveys by MacKenzie and Clement (2014, 2016) were optimised for estimating total population size, rather than dolphin distribution. Therefore, survey effort was low in areas where few dolphins were expected. To fill in these gaps, Roberts et al. (2019) developed a habitat model using environmental and biological variables to predict where dolphins are likely to be found in higher and lower densities.

Because Roberts et al. (2019) had access to very limited in-situ environmental data, they based the habitat model on the presence or absence of particular fish species (in a sparse set of inshore fish surveys conducted by NIWA), and satellite data on water turbidity by season.

Hector’s dolphins are often found in turbid water (e.g. Bräger et al. 2003). While turbidity could potentially act as a proxy for another environmental variable (e.g. prey availability), there is no evidence to suggest that turbidity *per se* has a direct effect. The hypothesis that turbidity is important to Hector’s dolphins does not explain how the species managed to flourish before large scale deforestation and intensive land use created current levels of turbidity.

Turbidity was favoured in the modelling process because the dolphin abundance data (MacKenzie and Clement 2014, 2016) had a seasonal signal. Hector’s dolphin distribution is more dispersed in winter than in summer, with respect to water depth and distance from land (Rayment et al. 2010; MacKenzie and Clement 2014, 2016). Turbidity was the only environmental variable available to the modelling team that had any seasonal component. Hence it was inevitable that the habitat model would suggest a correlation between dolphin distribution and turbidity. This is not an indication that turbidity drives dolphin distribution, but a demonstration that the modelling process was flawed. A better fit is likely to result from including depth and season (summer, autumn, winter, spring) directly in the model. Depth is *directly* relevant to Hector’s dolphins, because they feed mostly on bottom-dwelling fish (Miller et al. 2013).

There are also serious problems with the manner in which the statistical modelling has been done:

- It did not meet usual standards for model checking and diagnostics (e.g. Redfern et al. 2006, Elith and Leathwick 2007, Guillera-Arroita et al. 2015). For example, the standard practice of developing the model with part of the dataset and testing it with the remaining data was not followed.
- The fish data used are presence/absence only, with no accounting for differences in sampling method. Mid-water and epipelagic species are important to Hector’s dolphin (Miller et al. 2013), but were not sampled effectively in the data used in the habitat model.
- The best-fitting model was not chosen according to any quantitative and/or objective measure (e.g. Akaike’s Information Criterion). Instead, the model used was chosen at a workshop, based on opinions expressed by stakeholders. Evidence ratios (Anderson 2008), calculated directly from the AIC scores, show that the model chosen was 10.6 trillion times less likely than the best-fitting model.

These are critically important problems, especially in a model to estimate bycatch risk for different dolphin populations. Dolphin distributions predicted by the habitat model do not match observed data (Rayment et al. 2010, MacKenzie and Clement 2014). The Roberts et al. (2019) habitat model does not appear to be fit for purpose.

#### 3. Reproductive rate

The calculations of reproductive rate, and therefore how many dolphin deaths in fishing nets can be sustained, depend strongly on R_max_. Under contract to MPI, Edwards et al. (2018) adjusted the existing estimate of R_max_ upwards, using a curve of the relationship between body size and reproductive rate across a wide variety of mammals (Duncan et al 2007). There are five problems with this:

- The relationship illustrated in Duncan et al. (2007) reflects a very general emerging pattern that is not expected to hold under all circumstances. Exceptions are entirely valid.
- Hector’s dolphin would be *expected* to be an outlier. It is the world’s smallest dolphin, living in cool temperate waters in which calves must be large and well insulated to survive. Therefore, breeding females invest a large amount of energy into each calf, which is likely to reduce the rate at which they can produce calves.
- When a data point does not fit an expected relationship, it is not legitimate to simply move the data point. The normal science process involves fitting lines to data points, not data points to lines.
- There is no empirical evidence to support the revised-upwards estimate of R_max_ for Hector’s dolphins. Indeed the 2019-2020 field season produced the lowest ratio of calves to non-calves (0.8%) recorded since 1991 (Dawson and Slooten unpub. data).
- The use of a general relationship between body size and R_max_ is an appropriate last resort only for a species for which no relevant biological data are available. It is not an argument for ignoring existing biological data.

#### 4. Fishing effort data

The MPI approach requires fishing effort data with GPS locations. Despite GPS being widely available since the early 1990s, this level of precision has only recently been required from fishers by MPI (Figure 5). The quality of these location data has been questioned. A fishing industry representative taking part in the Expert Panel workshop in 2018 (Taylor et al. 2018) pointed out that the current data are not accurate, and do not meet the lower standard of being representative. For example, larger vessels are more likely to report the GPS locations of their fishing effort. Data on the spatial distribution of fishing effort are critical to estimating the risk of dolphin capture. The current data are apparently taken at face value by MPI.

**Figure 5.**
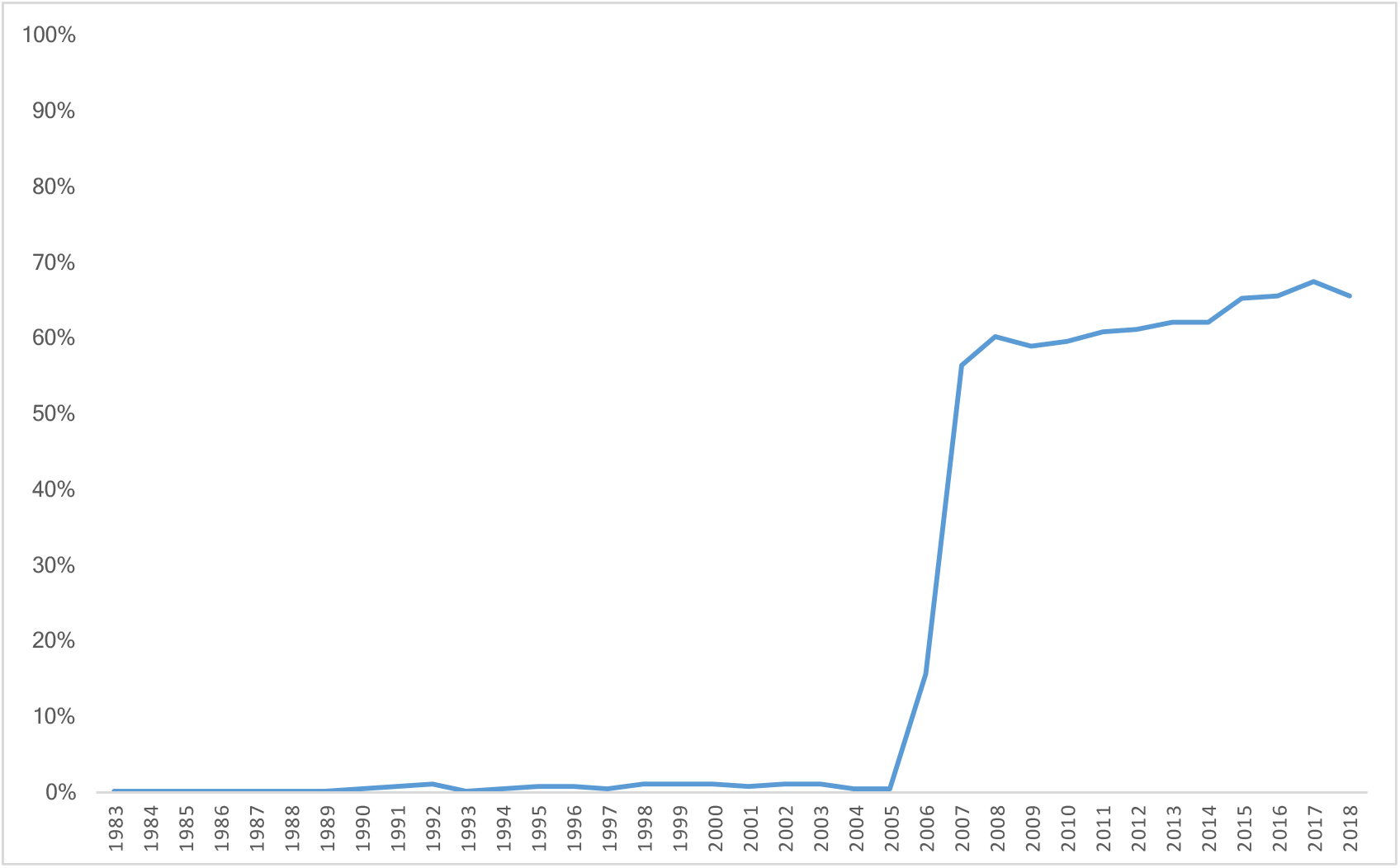
Proportion of gillnet fishing effort for which location is reported by latitude and longitude. Data provided by the Ministry for Primary Industries (MPI).

Until 2005, GPS locations were reported for less than 1% of the gillnet fishing effort. This increased to just over 15% in 2006, 56% in 2007, around 60% until 2015 and around 65% in recent years (Figure 5). The remaining fishing effort is reported by fisheries statistical areas (Figure 6). It is not clear how Roberts et al. (2019) created their maps of the spatial distribution of current fishing effort (from 65% of fishing effort currently reported with GPS locations). Likewise, it is not clear what assumptions were made in estimating the spatial distribution of past fishing effort, from current effort data.

**Figure 6.**
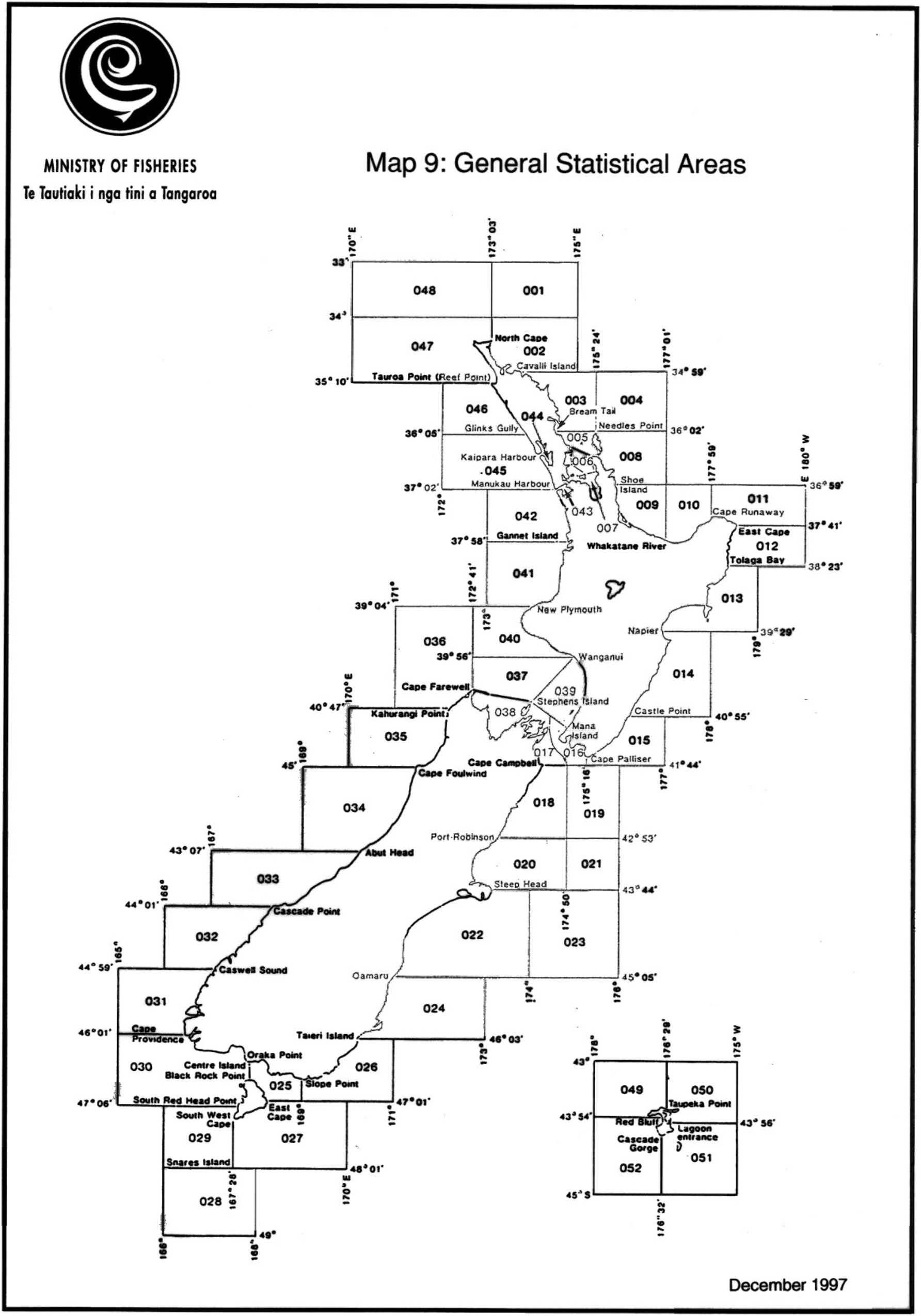
Fisheries statistical areas. Before 2006, the location of more than 99% of gillnet fishing effort was reported at the level of these statistical areas. In recent years, the location of about 35% of gillnet fishing effort is still reported at the level of these statistical areas (also see Figure 3).

The TMP estimate of current bycatch is 59 Hector’s dolphins per year (95% CI 23-127). Current gillnet fishing effort in Hector’s and Maui dolphin habitat is about 8,000 fishing days per year (Figure 7), compared to c. 20,000 days per year during the 1990s and 35,000 days per year in the early 1980s. If there had been no change in the size of the dolphin population, and no change in the spatial distribution of fishing effort, one would expect an average of about 150 dolphin deaths per year during the 1990s and early 2000s, and about 260 dolphin deaths per year during the early 1980s, based on the higher fishing effort alone. This is consistent with estimates from a team of government scientists and fishing industry experts(Davies et al. 2008), who estimated 110-150 Hector’s dolphin deaths per year during 2000-2006. Bycatch mortality of 260 Hector’s dolphins per year is on the order of 3 times the PST, and 23 times the PBR for the current dolphin population. Population size in the 1980s, and therefore bycatch, is likely to have been much higher than today, as estimated in previous risk assessments (e.g. Slooten and Dawson 2010, Davies et al. 2008).

**Figure 7.**
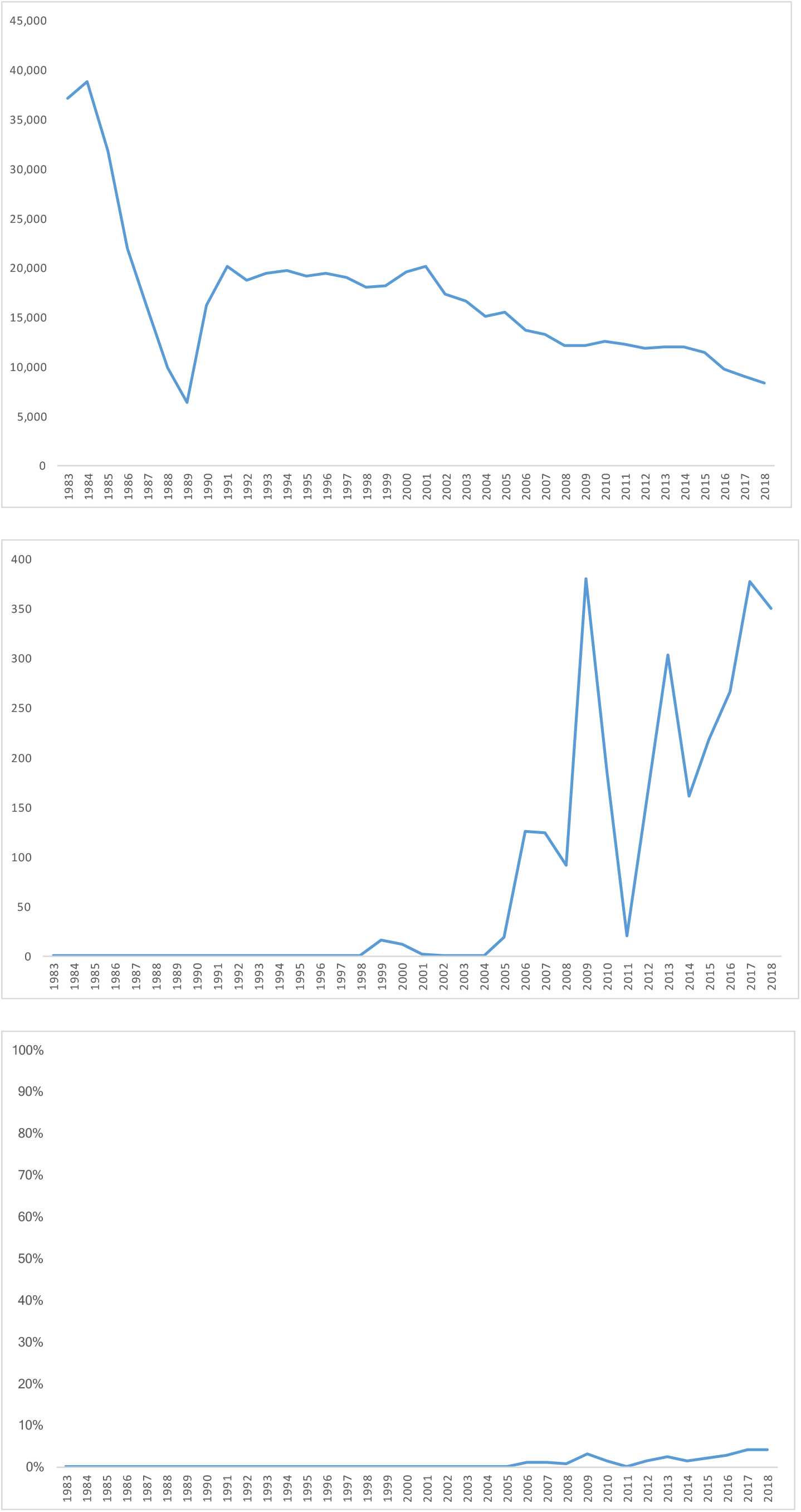
Gillnet fishing effort, in areas with Hector’s and Maui dolphin populations, in days (top), number of observed days (middle) and proportion of observed days (bottom). Data provided by the Ministry for Primary Industries (MPI).

In addition to much higher fishing effort in the 1980s and 1990s, the overlap between dolphins and gillnet fisheries was also much greater. Before 2008, the only areas with dolphin protection measures were Banks Peninsula and the North Island west coast. Since 2008, gillnets have been banned from the shoreline to 4 nautical miles offshore off most of the east and south coasts of the South Island.

#### 5. Observer coverage and estimates of bycatch

Observer coverage in New Zealand’s inshore fisheries is exceptionally low, and patchy in space and time (Figures 7–9). The only robust observer progamme for quantifying bycatch of Hector’s dolphins was achieved during the 1997/98 fishing season (Baird and Bradford 2000). Since then, observer coverage has been less than 5% in the gillnet (Figure 7), and inshore trawl fleet (Figure 8). The average number of observer days per year, in areas where Hector’s and Maui dolphins are found, increased briefly, after the last TMP (2007) and then dropped again to less than 5 observer days per year during 2011-2018 (Figure 9). The MPI estimate of the entanglement rate in trawl fisheries is based on a single observed capture.

**Figure 8.**
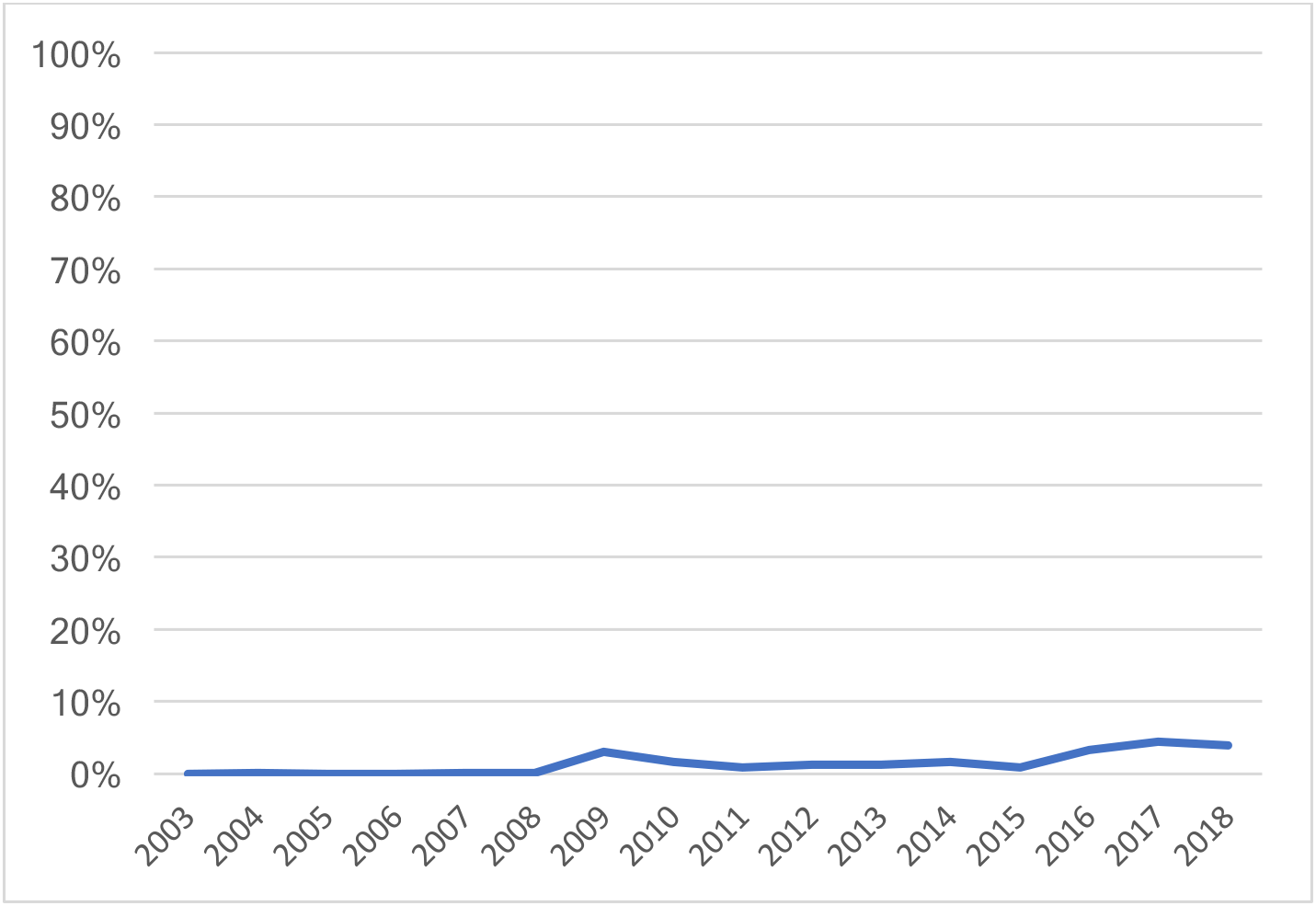
Proportion of observed trawling effort by inshore vessels (6-17 m in length) from 2003-2018. From Dragonfly (2020).

**Figure 9.**
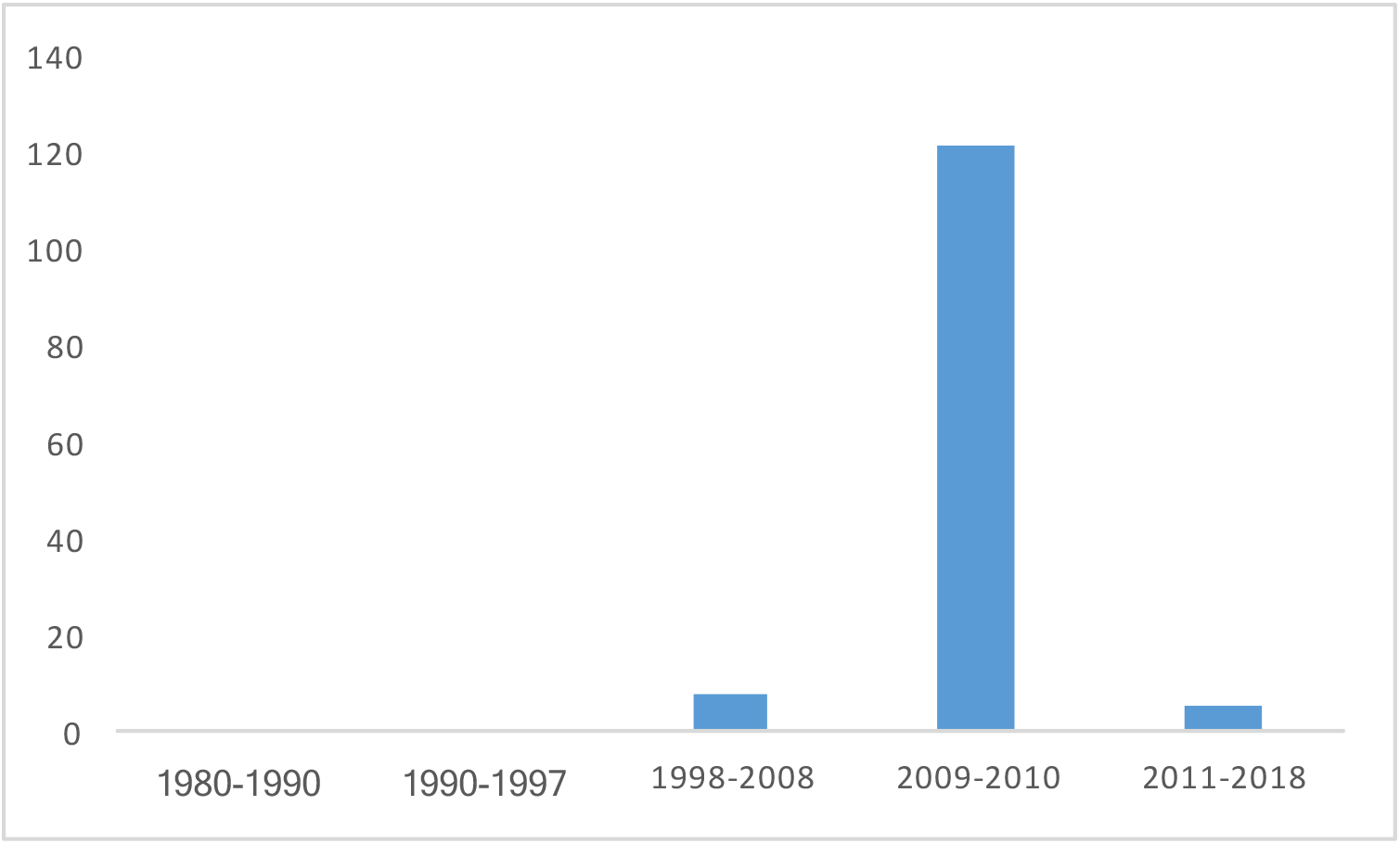
Average number of observer days per year, in the inshore gillnet fishery, in areas of Hector’s and Maui dolphin habitat. Data provided by the Ministry for Primary Industries (MPI).

The real problem is not that the bycatch estimates are highly uncertain, but that sampling theory indicates that they are likely to be biased low. For any one fishing vessel, catching a Hector’s dolphin is a relatively rare event. Low observer coverage acts to under-estimate bycatch.

We investigated this bias quantitatively via bootstrapping in R, for a gillnet fishery with 1000 gillnet sets in a season, 1000 years of monitoring and an underlying catch rate of 15 dolphins per 1000 gillnet sets (Figure 10). The fishing effort in the bootstrapping exercise is about double the size of the Timaru gillnet fishery in the 1997/98 observer programme. The catch rate used is lower than that found in the 1997/98 observer programme, but higher than that assumed by MPI to occur currently.

**Figure 10.**
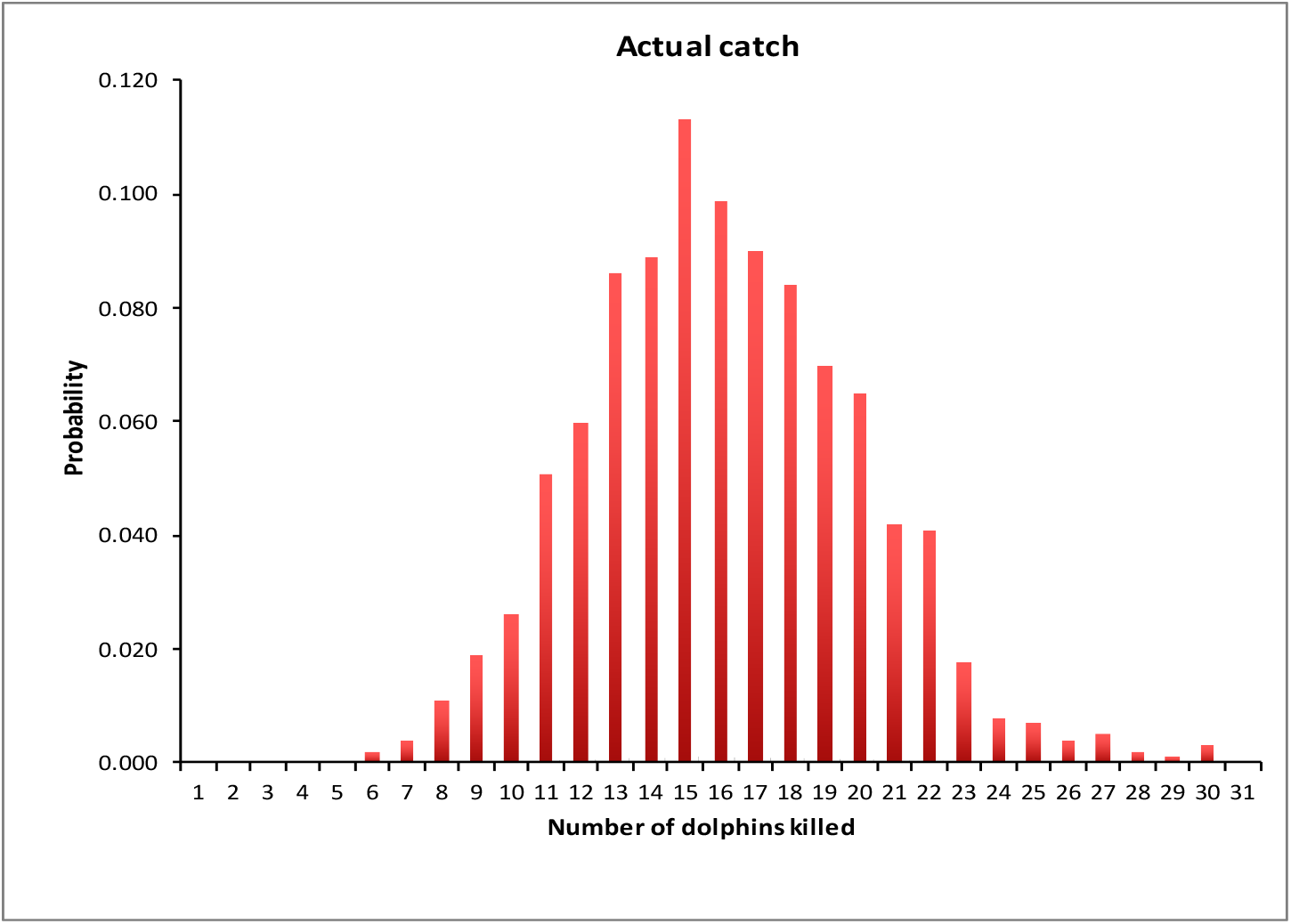
Bootstrap estimates of actual catch.

Notwithstanding significant levels of actual catch, at low (but typical) levels of observer coverage, the likelihood of observing a catch is also very low. With the 2% observer coverage in the bootstrapping exercise above, there is a 75% chance of seeing no captures in any one year (Figure 11).

**Figure 11.**
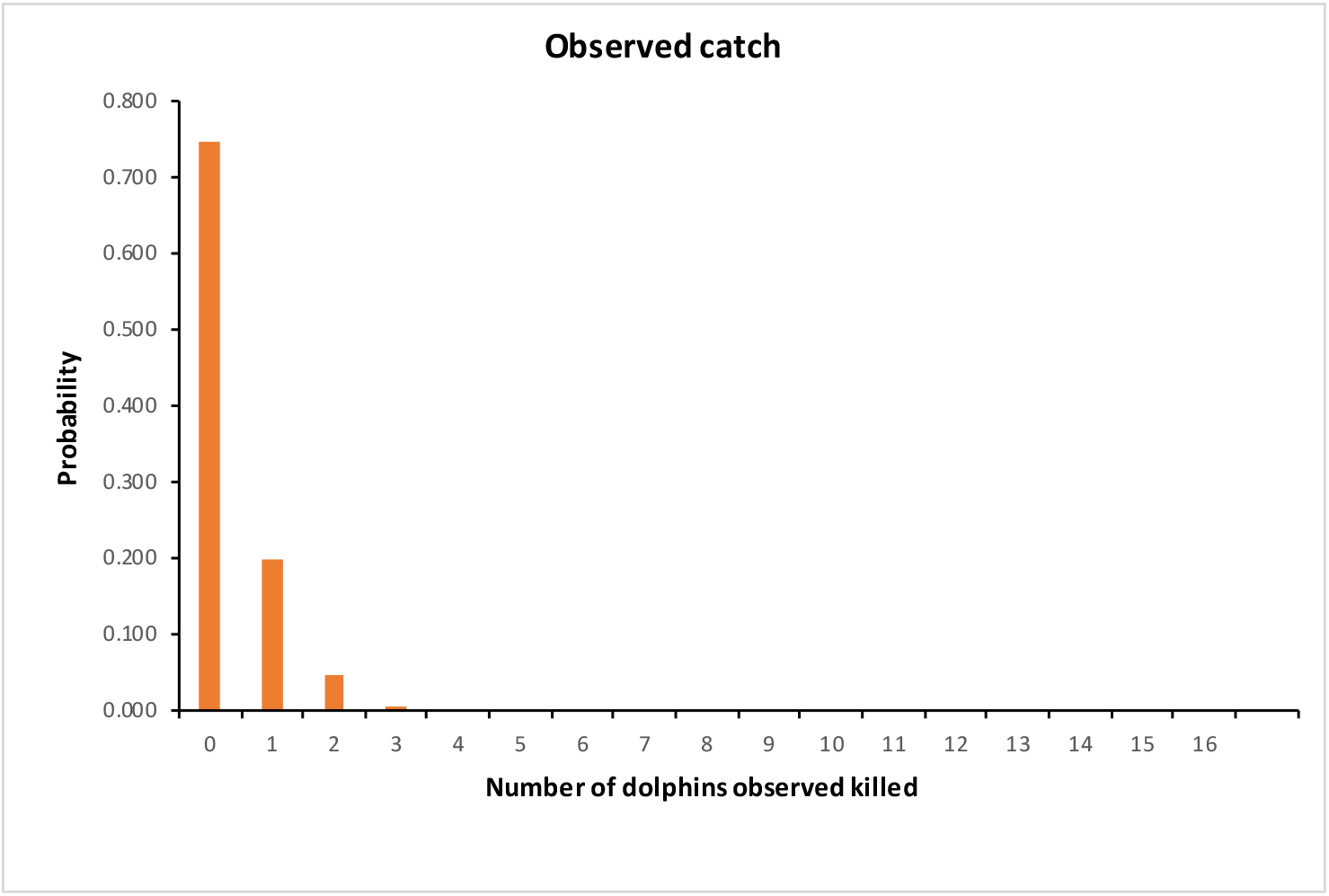
Observed catch at 2% observer coverage.

Extending these trials to cover a wider range of values for observer coverage and plausible bycatch rates shows that the very low level of recent observer coverage (1-3%) has resulted in estiimates of bycatch rate that are almost certainly biased low (Table 1).

**Table 1:** Probabilities of observing zero bycatch at given levels of observer coverage 1-10%) and true catch rate (0.4-4%). Hence a 1% catch rate combined with 2% observer coverage results in an 81.7% chance of observing zero bycatch in that observer programme. Note that these calculations assume that bycatch events are independent. If they are not, as could be caused by dolphins living in groups affecting the likelihood of multiple captures, probabilities of observing zero byatch would be higher than shown above.

| True catch rate | Observer coverage |  |  |  |  |
| --- | --- | --- | --- | --- | --- |
|  | 1.00% | 2.00% | 3.00% | 5.00% | 10.00% |
| 0.50% | 0.951 | 0.907 | 0.865 | 0.775 | 0.602 |
| 1.00% | 0.901 | 0.817 | 0.739 | 0.602 | 0.362 |
| 2.00% | 0.818 | 0.669 | 0.544 | 0.37 | 0.131 |
| 3.00% | 0.733 | 0.544 | 0.398 | 0.218 | 0.052 |
| 4.00% | 0.659 | 0.436 | 0.298 | 0.13 | 0.016 |

The US guidelines for preparing stock assessment reports (GAMMS 2016) address this problem specifically. In their terminology, the observer coverage applied in NZ inshore fisheries results in bycatch estimates that are “always biased”. This is a fundamental sampling problem that is not solved by the Bayesian approach taken in the analysis by Roberts et al. (2019).

That bycatch has been underestimated is consistent with Cooke et al.’s (2019) analysis of Maui dolphin population trends, in which the best fitting models used absolute levels of bycatch 15-20 times higher than those assumed by Roberts et al. (2019).

All observed catches have occurred at Kaikoura (area 18), north of Banks Peninsula (area 20) or south of Banks Peninsula down to Timaru (area 22). All of these areas have relatively high fishing effort. Average, reported fishing effort during 2015-2018 was 390 km of gillnet / year on the north side of Banks Peninsula (area 20) and 982 km / year on the south side of the peninsula to Timaru in the south (area 22). Most of the observed captures (13) have been in the Banks Peninsula area, which has relatively high fishing effort and high dolphin densities (thousands of Hector’s dolphins). A smaller number of captures (4) have been observed in Kaikoura (area 18). This area has the highest fishing effort recorded in any of the fisheries statistical areas, with 1,372 km of gillnets used / year during 2015-2018, but a relatively small population of a few hundred Hector’s dolphins. Areas where no catches have been observed either have little or no observer coverage, low fishing effort, small dolphin populations, or a combination of these factors.

In several areas, the timing of observer coverage has reduced the probability of detecting bycatch, with low or no observer coverage during years of high fishing effort. For example, area 34 (on the west coast of the South Island) saw high fishing effort (c. 1200 km / year) throughout the late 1980s and early 1990s. The only observer coverage in this area, however, has been 11 observer days in 2009 and 10 observer days in 2013, when fishing effort was below 100 km of gillnet per year. In most areas, there was no observer coverage until 2008, after the last TMP (2007) which substantially reduced the overlap between dolphins and gillnet fisheries (e.g. North Island west coast areas, Figure 12).

**Figure 12.**
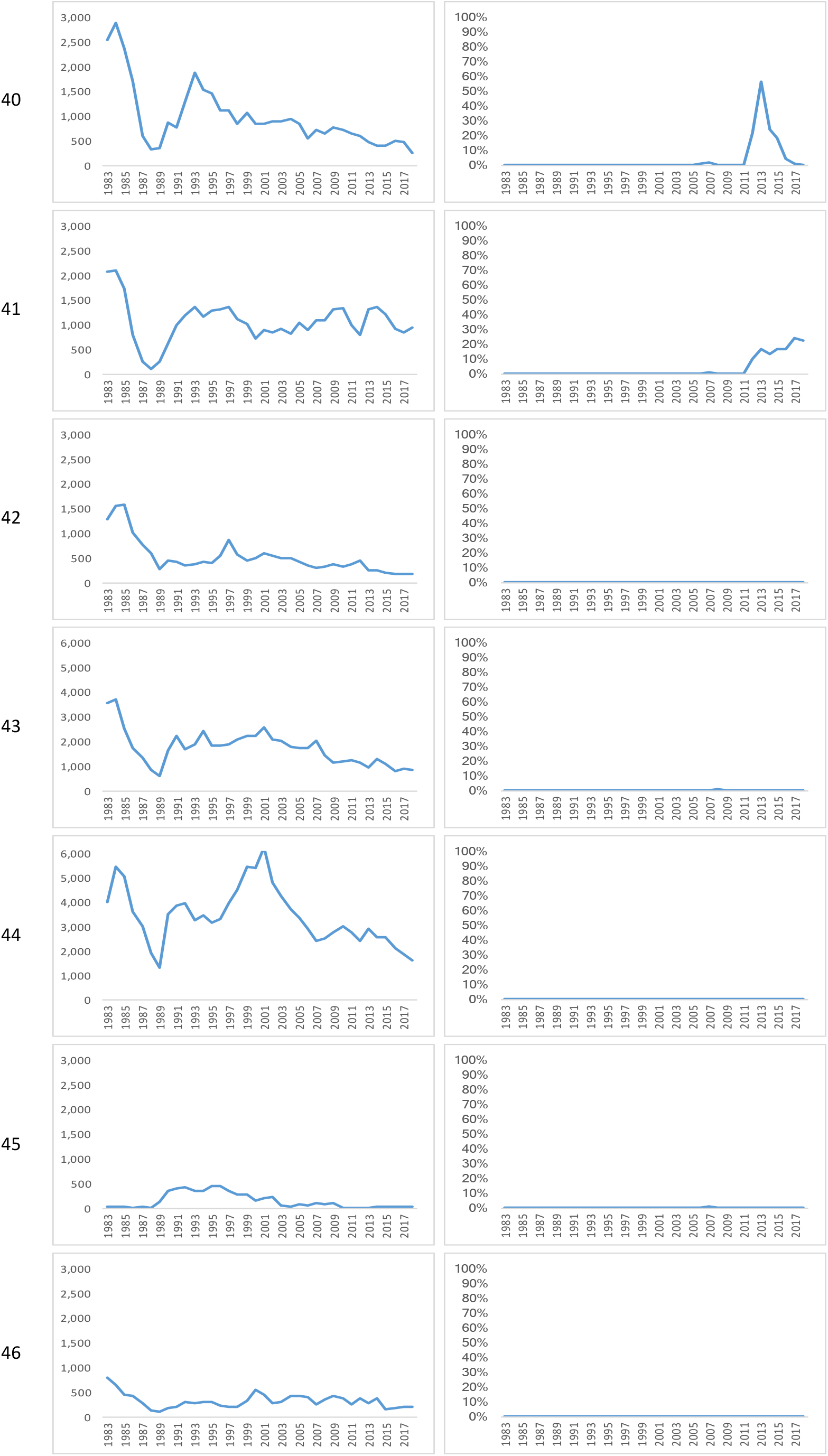
Gillnet fishing effort (left) and observer coverage (right) for fisheries statistical areas 40-46, off the west coast of the North Island. Figure 6 is a map showing the location of these fisheries areas. The y-axis for fishing effort, extends to 3,000 km of gillnet used per year for areas with relatively low fishing effort, and 6,000 km / year for areas with relatively high fishing effort.

Due to the problems illustrated in table 1, areas in which no bycatch has been observed are not necessarily low risk areas. They cannot safely be left out of the protection measures.

A substantial number of multiple catches, of several Hector’s dolphins caught in the same gillnet or trawl (Slooten et al. 2019), makes it even more difficult to obtain robust estimates of bycatch. MPI estimate that there are currently 59 Hector’s dolphins killed each year in commercial gillnet and trawl fisheries (95% ci 23-127; TMP 2019). Yet observer coverage has been so low that only 7 Hector’s dolphin catches have been observed in the last two decades.

The MPI habitat model takes bycatch estimates at face value, not recognising the high likelihood of bias. A properly designed observer programme is needed to obtain scientifically robust estimates of bycatch.

#### 6. PST - the metric used to determine a sustainable take

The Potential Biological Removal (PBR) method has been thoroughly tested via simulation (Wade 1998). MPI favours a NZ-developed method, known as the Population Sustainability Threshold (PST). The PST formula uses *ϕ* a so-called tuning parameter, that conflates uncertainty and the recovery objective. The PST approach has had limited peer review, and has not been published in a peer-reviewed scientific journal. It has not been subjected to the extensive testing via simulation that the PBR has, and it is not clear that the resulting estimates of acceptable bycatch are truly sustainable. That these estimates are far larger than PBR estimates of sustainable take is not reassuring (Table 2).

**Table 2.** Comparison of the PBR with the limit on fisheries mortality (PST) proposed by the New Zealand Ministry for Primary Industries (MPI) for the national, regional and local populations listed in the TMP.

|  | Estimated Bycatch | PBR | PST |
| --- | --- | --- | --- |
| **Hector's dolphin** | 59 | 11 | 74 |
| Regional populations of Hector's dolphin: |  |  |  |
| **East coast South Island** | 51 | 7 | 49 |
| **West coast South Island** | 6 | 4 | 27 |
| **South coast South Island** | 1 | 0.2 | 2 |
| **North coast South Island** | 1 | 0.1 | 1 |
| Local populations off east coast South Island: |  |  |  |
| **Kaikoura** | 11 | 1 | 10 |
| **Banks Peninsula** | 17 | 9 | 56 |
| **Timaru** | 20 | 3 | 34 |

The US approach was developed by marine mammal scientists who were deeply aware of meta-population theory, population fragmentation and source-sink dynamics. It therefore includes strategies to avoid depletion of local populations (Wade 1998). Acknowledgement of these issues as serious problems is absent in the TMP. For example, the PBR approiach requires limits to be calculated for populations that are demographically separate from others (i.e. a dolphin removed from one population will not likely be replaced from another population). The New Zealand PST approach includes none of these safeguards. The relative performance of the PBR and PST should be thoroughly tested, before the PST is used to inform management decisions.

#### 7. Estimating the incidence of non-fishery causes of death

Necropsies of dolphins that washed up on beaches (Roe et al. 2013) were used to estimate the number of Maui and Hector’s dolphin deaths from disease. This is a biased sample because most of the animals being dissected would be expected to be ill or old. As the current COVID-19 situation demonstrates, animals near the end of their lifespan are highly susceptible to disease. Basing conclusions on a beachcast sample exaggerates the real importance of disease, and cannot properly account for other contributing factors.

The expert panel (Taylor et al. 2018) concluded that *“we are not convinced that it is appropriate for the toxoplasmosis necropsy data to receive the full modelling treatment: the uncertainties and potential biases in these data are too large. If the effects of the disease are as large as they appear, and the deaths are additional to other causes, we would expect the populations of Hector’s dolphins to be in rapid free-fall towards extinction” (point 32, p.12)*

MPI’s estimates of the importance of disease are not consistent with published field observations:

- The survival rate of dolphins at Banks Peninsula increased significantly (Gormley et al. 2012) after gillnetting was banned inshore in 1988. If disease was an important problem this management change would have made negligible difference.
- The trend in genetic mark-recapture estimates of Maui dolphin also suggest a slower current rate of population decline than in the past (Baker et al. 2016). If toxoplasmosis was as severe a problem as the TMP claims, this population should be continuing to decline rapidly.
- No incidences of toxoplasmosis have been found in Hector’s or Maui’s dolphin since 2013.

Together, these data indicate that the treat of toxoplasmosis has been substantially overestimated. The discussion of the impact of disease in the TMP ignores the fact that it is, effectively, unmanageable. By contrast, bycatch is readily manageable.

## THE TMP’S SCIENTIFIC APPROACH: TWO OVERARCHING PROBLEMS

A. The first overarching problem is that the Roberts et al. (2019) approach combines several estimates that are biased, and the biases consistently act together to underestimate the level of bycatch and overestimate the species’ ability to absorb impacts. In essence, the approach uses abundance estimates that are likely biased high, multiplies them by a reproductive rate that has been arbitrarily raised, multiplied by an assumed figure for calf survival, to reach a number of dolphins that would be added each year if the population were to remain stable. From this number, Roberts et al. (2019) subtract their estimates of bycatch, which are almost certainly biased low. The remaining number of dolphins is apportioned a cause of death according to autopsy data from 55 Hector’s and Maui dolphins found dead on beaches. This is then compared to estimates of what level of takes would be sustainable, calculated using a formula that is not well understood and less conservative than the PBR. This is a poor basis for rational management of an endangered, endemic marine mammal.
B. The second overarching problem is that the Roberts et al. (2019) approach to “risk” does not relate to the risk of population decline or extinction, and is inconsistent with the modern understanding of the behaviour of meta-populations. The approach defines risk as the likelihood of capture, which is apportioned to different areas according to fishing effort and the habitat model’s outputs for dolphin distribution. The protection options are therefore targeted where high densities of dolphins and high fishing effort coincide. Large populations are allocated the highest level of protection, while small populations remain poorly protected. This approach makes no sense from a meta-population point of view. When fishing effort shifts from protected areas, to neighbouring, unprotected or poorly protected populations, small local populations are likely to decline. Population fragmentation is likely to increase. This approach would increase the risk of local extinctions, contractions of dolphin distribution, loss of genetic variability and result in increased risk to the species as a whole.

Many of these points were made in the Expert Panel workshop in 2018 (Taylor et al. 2018). Despite repeated requests (e.g. at stakeholder meetings) MPI have failed to provide a list of recommendations from the Expert Panel Report with their responses (if any) to those recommendations. This standard step in scientific practice, following peer review (e.g. response to reviewers’ comments) has not been followed.

